# Mouse nephron formation is impaired by moderate-dose arsenical exposure

**DOI:** 10.1101/2024.09.06.611740

**Authors:** Carlos Agustin Isidro Alonso, Jenna Haverfield, Gabriela Regalado, Sihem Sellami, Natascha Gagnon, Ajay Rajaram, Pierre Olivier Fiset, Aimee K Ryan, Koren K Mann, Indra R Gupta

## Abstract

**Background:** Arsenic is a naturally occurring toxicant and industrial byproduct with significant health risks. Globally, millions of people are exposed to arsenic concentrations that exceed the World Health Organization’s recommended limit of 10 μg/L. Chronic arsenic exposure is linked to an increased risk of chronic kidney disease (CKD); however, the effects of arsenic exposure on kidney development remain unclear. Eukaryotes methylate inorganic arsenic (iAsIII) using the enzyme arsenic 3 methyltransferase (As3mt), that converts it to methylated intermediates, mono and dimethyl arsonous acid (MMAIII and DMAIII), and mono and dimethyl arsonic acid (MMAV and DMAV). We hypothesized that arsenicals exposure during mouse kidney development impairs nephron formation.

**Methods:** Cultured mouse embryonic kidney explants were treated with inorganic arsenite (iAsIII), MMAIII, MMAV, and DMAV. Female mice harboring a humanized version of *AS3MT* and wild-type mice with murine *As3mt* were exposed to iAsIII throughout gestation and weaning and their offspring were analyzed for kidney defects.

**Results:** Inorganic arsenic, iAsIII (200 μg/L), inhibited ureteric bud branching morphogenesis and growth of mouse kidneys at embryonic day 11.5 (E11.5) and E12.5, but not at E13.5. MMAIII, but not MMAV or DMAV, impaired ureteric bud branching and kidney explant growth. Additionally, iAsIII exposure increased apoptosis in the metanephric mesenchyme of E11.5 explants and decreased *Gdnf* transcription, which may explain the impairment in ureteric bud branching. Humanized mouse pups exposed to 200 *μg/L* iAsIII *in utero*, showed a 20% reduction in kidney weight normalized to body weight and a 28% reduction in nephron number, compared to kidneys of wild-type mice.

**Conclusion:** Exposure to arsenicals during embryonic development impairs ureteric bud branching morphogenesis and decreases nephron endowment, which may predispose to CKD in adulthood.

## Introduction

Inorganic arsenic (iAs) is a toxic metalloid and one of the top 10 environmental toxicants of concern listed by the World Health Organization (WHO). As a natural component of the earth’s crust, iAs leaches into groundwater that may be used for consumption and crop irrigation. Private well water is the primary source of iAs exposure globally, however, industrial emissions arising as byproducts of smelting, hydraulic fracturing and mining are other sources. It is estimated that 2.1 M people in the US depend on well water contaminated with levels of iAs greater than the WHO’s recommended maximum concentration of 10 μg/L or 10 ppb in their drinking water and this number ascends to 140 million people worldwide ^1–4^.

iAs is bio-transformed through methylation by the enzyme arsenite methyltransferase, As3MT(GeneID Human: 57412; Mouse:57344). Methylation of iAs produces monomethylated (MMAs) and dimethylated (DMAs) arsenicals that exist in trivalent or pentavalent state. While iAs methylation has traditionally been considered a detoxification process, some methylated intermediates may be more toxic than iAs, with the trivalent forms generally being more cyto- and geno-toxic than the pentavalent forms ^5,6^. Acute poisoning with high dose iAs from industrial exposures and through suicide attempts results in acute kidney failure and acute tubulointerstitial nephritis ^7^. In humans, chronic exposure of moderate to high concentrations, greater than 100 μg/L, may also be nephrotoxic, resulting in microalbuminuria, proteinuria, and impaired kidney function as measured by serum creatinine levels. The susceptibility of the developing kidney to iAs-induced nephrotoxicity is not known ^8^.

Malformations in human kidney arise during human gestation beginning as early as four weeks of gestation and continuing until birth. Severe kidney malformations include failure to form a kidney(s) or formation of a small, dysplastic or hypoplastic kidney(s). Indeed, these types of malformations are the most common cause of kidney failure in childhood and in adults up to middle age ^9^. Analysis of genetically modified mouse models show that loss-of-function mutations in specific genes can result in a spectrum of phenotypes, from gross kidney malformations to more subtle defects like a marked reduction in nephron number ^10–13^. Although at least 54 genes are implicated in monogenic kidney malformations in humans ^14^, less is known about the environmental factors contributing to kidney malformations and/or defects in nephrogenesis during fetal development ^15^. Indeed, the impact of maternal exposure to environmental toxicants and fetal kidney development has been largely unexplored.

We hypothesized that exposure to arsenicals is toxic to the embryonic kidney and may impair nephron formation. To address this hypothesis, we present two models of exposure: *ex vivo* mouse embryonic kidney explants, and *in vivo* pregnant transgenic mice harboring a humanized version of the gene encoding the arsenic metabolizing enzyme, As3MT.

## Materials and Methods

### Animal models and exposures

All procedures involving mice received ethics approval by the McGill University Animal Care Committee (Animal Use Protocol numbers MUHC-4120, JGH-5664, JGH-10000) in accordance with the Canadian Council on Animal Care guidelines.

For the *ex vivo* experiments, *Hoxb7-GFP* females (MGI:4946625) ^16^(Srinivas et al., 1999) were mated with WT CD1 males. Females were checked for plugs daily and separated when a plug was found (E0.5). Pregnant females were euthanized at days E11.5, E12.5 or E13.5 and *Hoxb7-GFP* positive embryos were dissected to obtain embryonic kidneys. For the *in vivo* experiments, C57BL/6J mice harboring the mouse *As3mt* locus (termed ‘wild-type’ mice) were purchased (Jackson Laboratory). C57BL/6J mice with the human *AS3MT locus* (termed ‘humanized’ mice) ^17^ were obtained from the Mutant Mouse Resource and Research Centre. Two weeks prior to mating, all mice were fed AIN-76 chow (Envigo, Lachine, Quebec), which contains low levels of iAs ^18,19^. Pregnant dams were exposed to tap water with iAs levels below 0.05 μg/L, or tap water supplemented with 200 μg/L NaAsO_2_ (iAs(III)) throughout gestation until postnatal day 14 (P14), when nephrogenesis ends in mice. iAsIII-containing drinking water was changed three times per week. At P14, the pups were weighed, euthanized and their kidneys were dissected and weighed.

### Embryonic kidney explant culture

Each pair of kidneys/embryo was dissected and placed on polyethylene terephthalate membrane inserts with 0.4 μm pores (Corning™ Falcon™ Cell Culture Inserts). One kidney was placed on an insert submerged in media (DMEM-F12 (Cat#319-085-CL Wisent) with 5% fetal bovine serum (Cat#080-150 Wisent) and 1% penicillin/streptomycin with water vehicle (Cat#A5955 Sigma),), while the mate kidney was submerged in media supplemented with one of the following arsenicals: 20 μg/L NaAsO_2_ (Millipore-Sigma, S7400), 200 μg/L NaAsO_2_, 200 μg/L monomethylarsonous acid (MMAIII), 200 μg/L monomethylarsonic acid (MMA V), or 200 μg/L dimethylarsonic acid (DMAV). MMAIII, MMAV and DMAV were generously gifted by Dr. D. Scott Bohle (Department of Chemistry, McGill University). Explants were cultured for 48 hours in a humidified incubator with 5% CO_2_. Epifluorescence and brightfield images were taken at 0, 24 and 48 h.

### Nephron number enumeration

Quantification of glomerular number per kidney at P14 was performed using the acid maceration technique ^20,21^. Two hundred μl of each kidney suspension was added to a well containing a 16-square grid on the base. The total number of glomeruli in each well was counted in triplicate using brightfield microscopy and used to derive a mean glomerular count.

### Arsenic speciation analysis

Arsenic speciation was performed on media samples to determine the proportion of inorganic arsenic, MMA and DMA, but was unable to distinguish between +3 and +5 valence states. Briefly, media samples were collected at the end of the embryonic kidney explant culture, snap frozen and stored at -80°C. Arsenic speciation for each sample was performed at the Dartmouth Trace Element Analysis Core (USA) using an anion exchange chromatography (50 mm Hamilton PRPX-100) with a 50 mM (NH_4_)_2_CO_3_ gradient on the Agilent™ 8900 triple quadrupole Inductive Coupled Plasma Mass Spectrometer coupled to an Agilent™ 1260 Liquid Chromatography system (Jackson et al., 2015). Detection limits for arsenic species are 10 ng/L.

### RNAScope™

RNA *in situ* hybridization was performed on 4% paraformaldehyde-fixed P14 kidney sections from wild-type and humanized mice using RNAscope™ (Advanced Cell Diagnostics). A mouse-specific *As3mt* probe (RNAscope™ Probe-Mm-As3mt-C2; 1299261-C2) was used for the wild-type mouse kidney sections, while a human-specific *AS3MT* probe (RNAscope™ Probe-Hs-AS3MT-C2; 518841-C2) was used for the humanized mouse kidney sections. The experimental procedure followed the protocol provided by Advanced Cell Diagnostics RNAscope™ Multiplex Fluorescent Reagent Kit v2 user manual. The sections were counterstained with Phalloidin-488 and DAPI to visualize tissue structure. Positive and negative control probes were included in each assay to validate the specificity and sensitivity of the staining. Representative areas of each slide were imaged using a Zeiss LSM880 Laser Scanning Confocal microscope.

### RT-qPCR

After 48 hours of culture, six E11.5 explants per treatment group were snap frozen. RNA was isolated using RNeasy Mini Kit (Qiagen 74104) following the manufacturer’s instructions. The purity and concentration of the RNA samples were evaluated using a NanoDrop One spectrophotometer (ThermoFisher Scientific). RNA was reverse transcribed using Moloney murine leukemia virus reverse transcriptase and random hexamer primers. qPCR reactions were performed using Blastaq Master mix Applied Biological Materials G891) using a LightCycler® 96 Instrument (Roche). Expression of *Rpl13* (GeneID: 6137), *As3mt* (57344), *Gdnf* (14573), *Ret* (19713), *Six2* (20472), and *Wt1* (22431) was analyzed in duplicate. Gene expression was determined relative to that of the reference gene Rpl13 using the 2-ΔΔCt method. Primer sets are listed in Table 1.

**Table 1:**
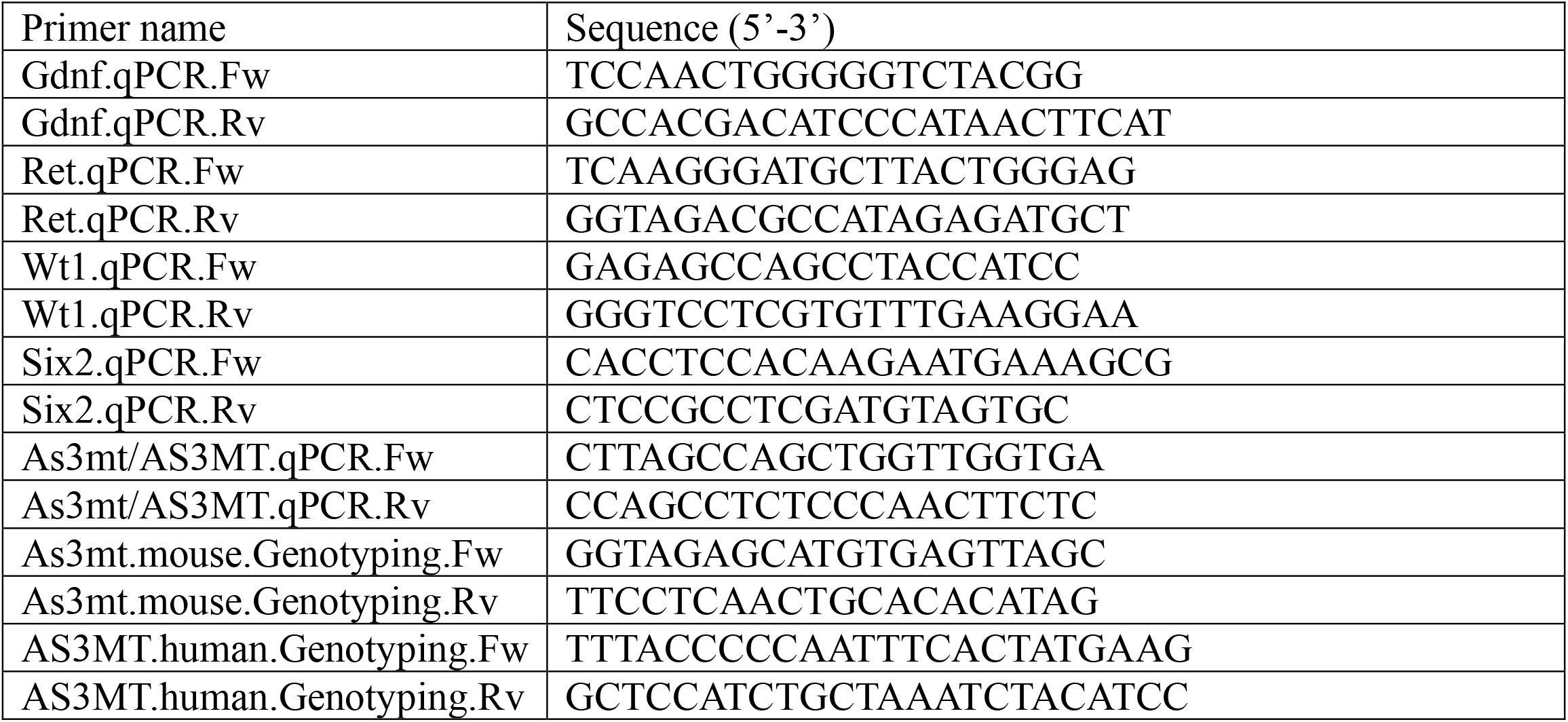
Primers list.

### Immunofluorescence and TUNEL staining

Embryonic kidney explants cultured for 48h were fixed with PFA 4% at 4°C overnight. After washing with PBS, explants were embedded in histoprep gel (Eprepin A11122) and processed, paraffin-embedded, and sectioned at 5 μm thickness. Sections were deparaffinized, rehydrated, and subjected to heat-induced antigen retrieval using EDTA buffer (1.2 mM EDTA pH8). After cooling, sections were blocked (10% normal goat serum) for 1 h. and incubated with anti-GFP primary antibody (Invitrogen, A11122; 1:100) overnight at 4°C. After washing, slides were incubated with an anti-rabbit IgG coupled to Alexa-488 (Invitrogen A11034) for 1 h at RT. After secondary antibody incubation, slides were washed and incubated with In Situ Cell Death Detection Kit, TMR red (Roche 12156792910) following the manufacturer’s instructions. Sections were counterstained with DAPI, and mounted with Fluorsave (Millipore Sigma 345789) mounting medium.

For whole mount immunofluorescence, kidney explants were fixed in 100% ice-cold methanol for 10 minutes, washed with PBST (PBS with 0.1% Tween 20), and blocked with 2.5% goat serum and 5% BSA in PBST overnight. Explants were then incubated with anti-SIX2 (Thermo Fisher 11562-1-AP; 1:100) and anti-WT1 (Santa Cruz sc7385; 1:100) primary antibodies overnight at 4°C. After washing, explants were incubated with secondary antibodies anti-rabbit IgG coupled to Alexa488 (Invitrogen A11034; 1:200) and anti-mouse IgG Alexa555 (Invitrogen A21126; 1:200) for 3 h at RT following by washing and counterstained with DAPI 1 μg/mL. Explants were placed on imaging dishes submerged on Fluorsave mounting media.

### Statistical analysis

Normality and equal variance were tested in all cases. For ureteric bud branching analysis, paired t-tests were employed. For kidney explant perimeter analysis, two-way ANOVA followed by a Holm-Sidak multiple comparison test was performed. For TUNEL, RT-qPCR. RNA-scope quantification, wild-type and humanized kidney weights, normalized wild-type kidney weight, and wild-type nephron endowment parametric t-tests were used. Normalized humanized kidney weight (control) and humanized nephron endowment (control) data did not pass the normality test (D’Agostino and Pearson test), thus the non-parametric Mann-Whitney t-test was used.

## Results

Nephrogenesis is defined by the iterative process of ureteric bud branching such that each ureteric bud tip induces the adjacent mesenchyme to form a nephron. The transgenic *Hoxb7-GFP* mouse line undergoes normal kidney development that can be visualized using green fluorescent protein (GFP) expressed in the ureteric bud and its derivatives throughout ureteric bud branching morphogenesis ^16^. To define the effects of arsenical exposure on ureteric bud branching, females carrying a *Hoxb7-GFP* allele were bred to wild-type males and the dams were euthanized at E11.5, E12.5 or E13.5. At each timepoint, the kidneys were dissected and grown as explants in culture with or without 200 μg/L (∼1.5 μM) sodium arsenite (iAsIII)-supplemented media to model an exposure to iAs at the beginning of ureteric bud branching.

*Exposure to inorganic arsenic and MMA-III impairs ureteric bud branching and kidney growth* A marked reduction in ureteric bud branching was observed in E11.5 and E12.5 kidney explants cultured for 48 h with 200 μg/L iAsIII, when compared to control explants (mean branching index); E11.5: control=3.78, iAsIII=0.67; E12.5: control=3.18, iAsIII=1.40, p<0.0001) (**Fig. 1A, 1C, 1D, 1F**). In contrast, no differences in branching were noted when E13.5 explants were cultured for 48 h with iAsIII (p=0.0857) **(Fig. 1G. 1I)**. Total kidney perimeter was significantly reduced in E11.5 explants cultured with 200 μg/L iAsIII after 24 h (control=1700μm, iAsIII=1367μm) and 48 h (control=2381μm, iAsIII=1314μm) of culture (p<0.0001)(**Fig. 1B**). After 48 h in culture, iAsIII-treated E12.5 explants also showed a significant reduction in explant perimeter (control=2745μm, iAsIII=2060μm, p<0.0001) (**Fig. 1E**). In contrast, iAsIII-treated E13.5 explants did not show changes in total kidney perimeter compared to control explants (p=0.8217). Explants cultured in the presence of iAsIII also exhibited a much darker cytoplasm (Supplemental **Fig. 2**). To assess iAsIII exposure at lower concentrations, E11.5 kidney explants were cultured with 20 μg/L iAsIII. No differences were detected in the branching index nor the kidney perimeter at this lower concentration (**Supplemental Fig. 1**).

**Fig 1.**
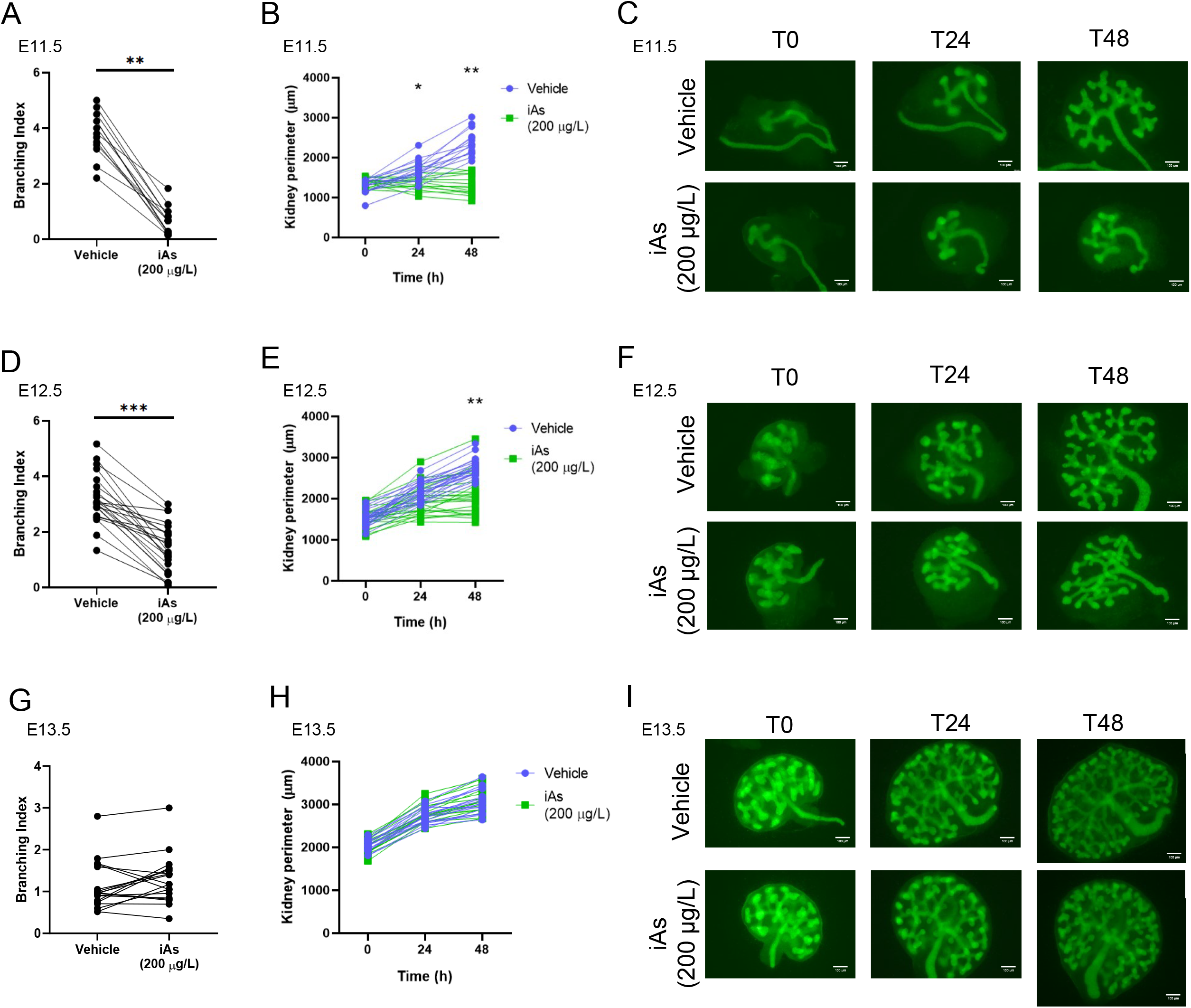
iAsIII impairs early ureteric bud branching and kidney growth. Embryonic kidneys were dissected from mouse embryos at embryonic day (E)11.5 (A, B, C), E12.5 (D, E, F) and E13.5 (G, H, I) and cultured in the presence or absence of 200 μg/L iAs. Ureteric branching index (A, D, G) was calculated as (N_T48_-N_T0_)/N_T0_, where N=ureteric bud tips. Data were analyzed using paired t-tests. Kidney perimeter (B, E, H) was measured at each timepoint from brightfield images (Supplemental Fig. 1). Data were analyzed using two-way ANOVA. Representative epifluorescence images (C, F, I) at T0, T24 and T48h of culture are shown. Scale bar equals 100 μm. A, B, C: n=17; D, E, F: n=23; G, H, I: n=19. Embryos are collected from at least 3 dams. * = p < 0.05; ** = p < 0.01; *** = p < 0.001.

**Fig 2.**
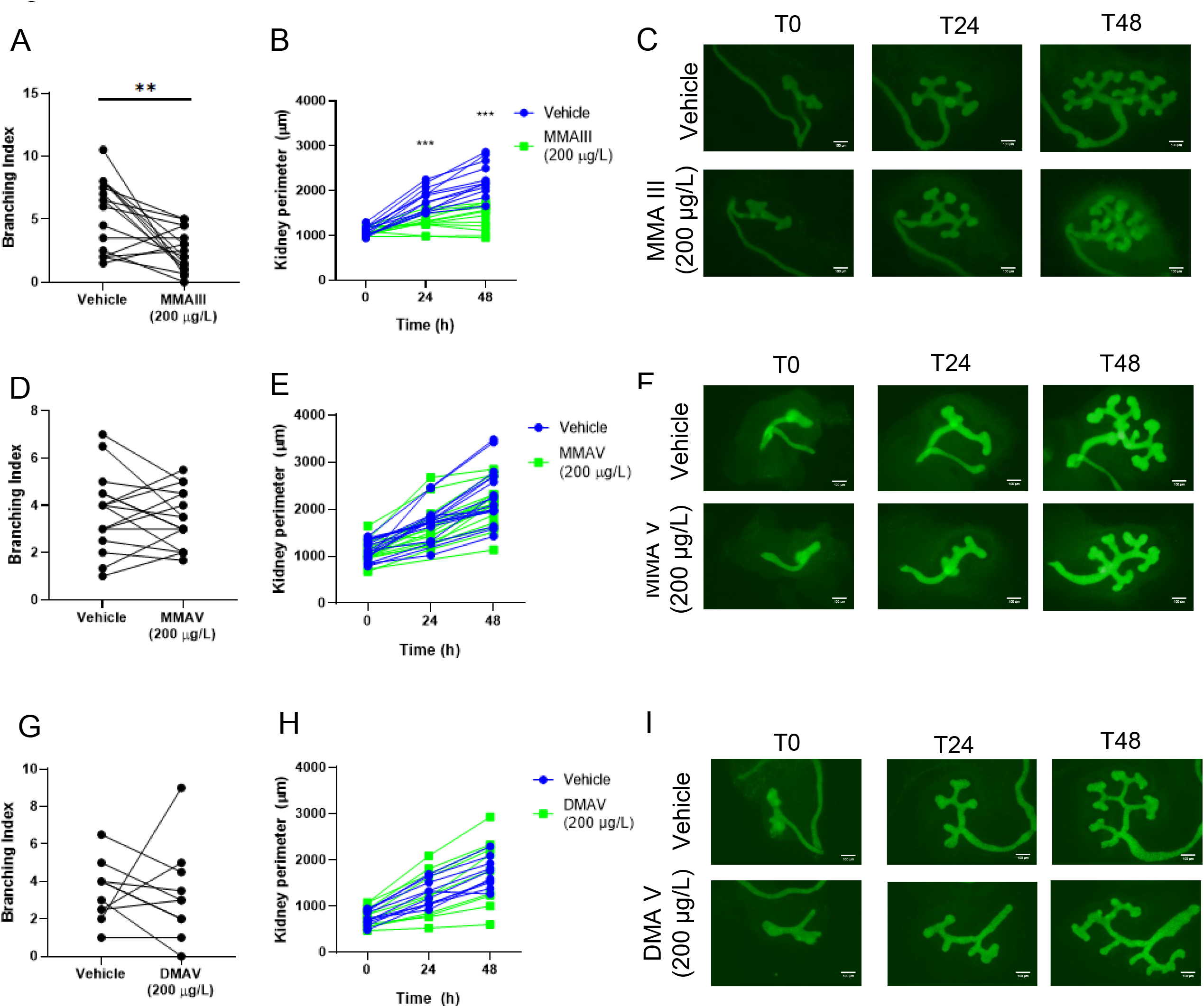
MMAIII, but not MMAV or DMAV, impairs ureteric branching and kidney growth. Embryonic kidneys were dissected from mouse embryos at E11.5 and cultured in the presence or absence of 200 μg/L MMAIII (A, B, C), MMAV (D, E, F) or DMAV (G, H, I). Ureteric branching index (A, D, G) was calculated as (N_T48_-N_T0_)/N_T0_. Data were analyzed using paired t-tests. Kidney perimeter (B, E, H) was measured at each timepoint from brightfield images (Supplemental Fig. 1). Data were analyzed using two-way ANOVA. Representative epifluorescence images (C, F, I) at T0, T24 and T48h of culture are shown. Scale bar equals 100 μm. A, B, C: n=17; D, E, F: n=16; G, H, I: n=11 from at least 3 dams. * = p < 0.05; ** = p < 0.01; *** = p < 0.001.

To test the toxicity of arsenic metabolites, E11.5 embryonic mouse kidney explants were exposed to 200 μg/L of MMAIII, MMAV, DMAV, or vehicle for 48 h (**Fig. 2**). Exposure to DMAIII was not assessed because the compound rapidly oxidizes to DMA-V in solution. Ureteric bud branching (control=5.53, MMMAIII=2.41, p=0.0016) and perimeter (control=2264μm, iAsIII=1408μm, p<0.0001 after 48 h were severely impaired in the presence of MMAIII at 200 μg/L (**Fig. 2A-C**). Similar to iAsIII, the MMAIII-treated explants exhibited a darker cytoplasm compared to the vehicle-treated (Supplemental **Fig. 2**). MMAV or DMAV (200 μg/L) did not cause changes in ureteric bud branching, kidney perimeter or aspect (**Fig. 2D-I, Supplemental Fig. 2**).

Since exposure to iAsIII (200 μg/L) and MMAIII (200 μg/L) resulted in smaller kidneys with less ureteric bud branching, cell survival was examined using the TUNEL assay. There was a marked increase in apoptotic cells, predominantly in the metanephric mesenchyme of iAsIII-treated explants compared to vehicle at E11.5 (mean number of apoptotic cells per tissue section area in μm^2^; control=696, iAsIII=3492, p=0.002) (**Fig. 3A-B**). GFP-positive tubules showed few to no apoptotic cells. To understand the molecular mechanisms underpinning the ureteric bud branching defects, we evaluated the expression of four genes involved in branching morphogenesis, *Gdnf, Ret, Wt1* and *Six2*, in vehicle and iAsIII-treated E.11.5 kidney explants. Interestingly, RT-qPCR revealed that *Gdnf*, which is expressed by the metanephric mesenchyme, was downregulated (46% reduction, p=0.033) following exposure to iAsIII, while the level of *Ret*, which is expressed by the ureteric bud epithelium, was not affected (p=0.23) (**Fig. 3C**). *Six2* and *Wt1* mRNA expression did not differ between vehicle and iAsIII-treated explants (**Fig. 3C**), which is consistent with our immunofluorescence results (Supplemental **Fig. 3**). The localization of SIX2 and WT1 were unaffected by iAsIII treatment (Six2: p=0.34 WT1: p=0.65) (**Supplemental Fig. 3**). GDNF and RET protein expression and localization were not assessed due to the lack of reliable antibodies.

**Fig 3.**
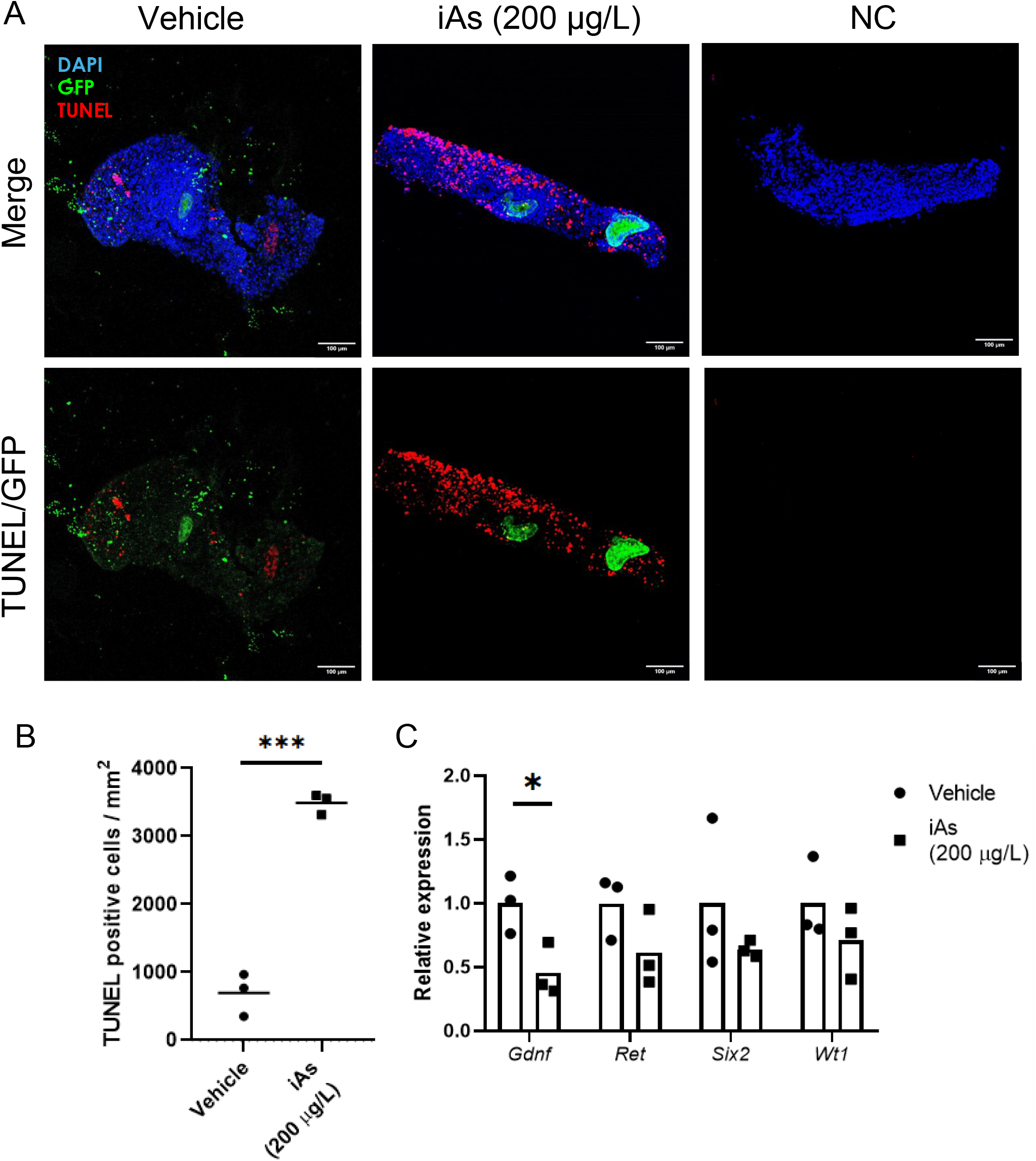
iAs increases apoptosis in the metanephric mesenchyme of embryonic kidney explants. E11.5 mouse kidneys were dissected and cultured for 48 h in the presence or absence of 200 μg/L iAs. Kidney explants were fixed, embedded and sectioned for immunofluorescent studies using anti-GFP (green) and a TUNEL (red) assay. Nuclei were counterstained with DAPI (A). Cells undergoing apoptosis were quantified (B) and normalized to explant area excluding GFP-positive tubules. RT-qPCR was performed on vehicle versus iAsIII-treated explants after 48 h of culture to assess molecular markers of ureteric bud branching morphogenesis. There was a significant downregulation of *Gdnf* mRNA compared to controls (C). n= 3. Data were analyzed using a t-test. *** = p < 0.001.

To confirm whether iAsIII is metabolized in mouse embryonic kidney explants, mRNA expression of *As3mt in situ* was examined, and arsenic speciation was performed on the culture media. *As3mt* transcripts were observed in the ureteric bud and in the surrounding metanephric mesenchyme at E11.5 of both vehicle- and iAsIII-treated kidney explants, suggesting that iAsIII did not affect the spatial distribution or level of *As3mt* expression (**Fig. 4A**). Additionally, RT-qPCR analysis of E11.5 kidney explants cultured for 48 h confirmed that there was no difference in *As3mt* expression levels (p=0.9595) (**Fig. 4B**). Culture media was analyzed for arsenic speciation after 48 h and revealed high amounts of iAs in the iAsIII-treated samples at E11.5 and E12.5, 163.4 and 170.1 μg/L, respectively (**Fig. 4C**). MMA was not detected in any of the aliquots, consistent with previous observations that mice are more efficient at metabolizing iAsIII to DMAs than humans ^22^. Indeed, small amounts of DMA were detected in the media aliquots from iAsIII-treated explants, confirming that the mouse embryonic kidney explants used in this study can methylate iAsIII. As3MT protein expression was not examined due to the lack of a specific antibody.

**Fig 4.**
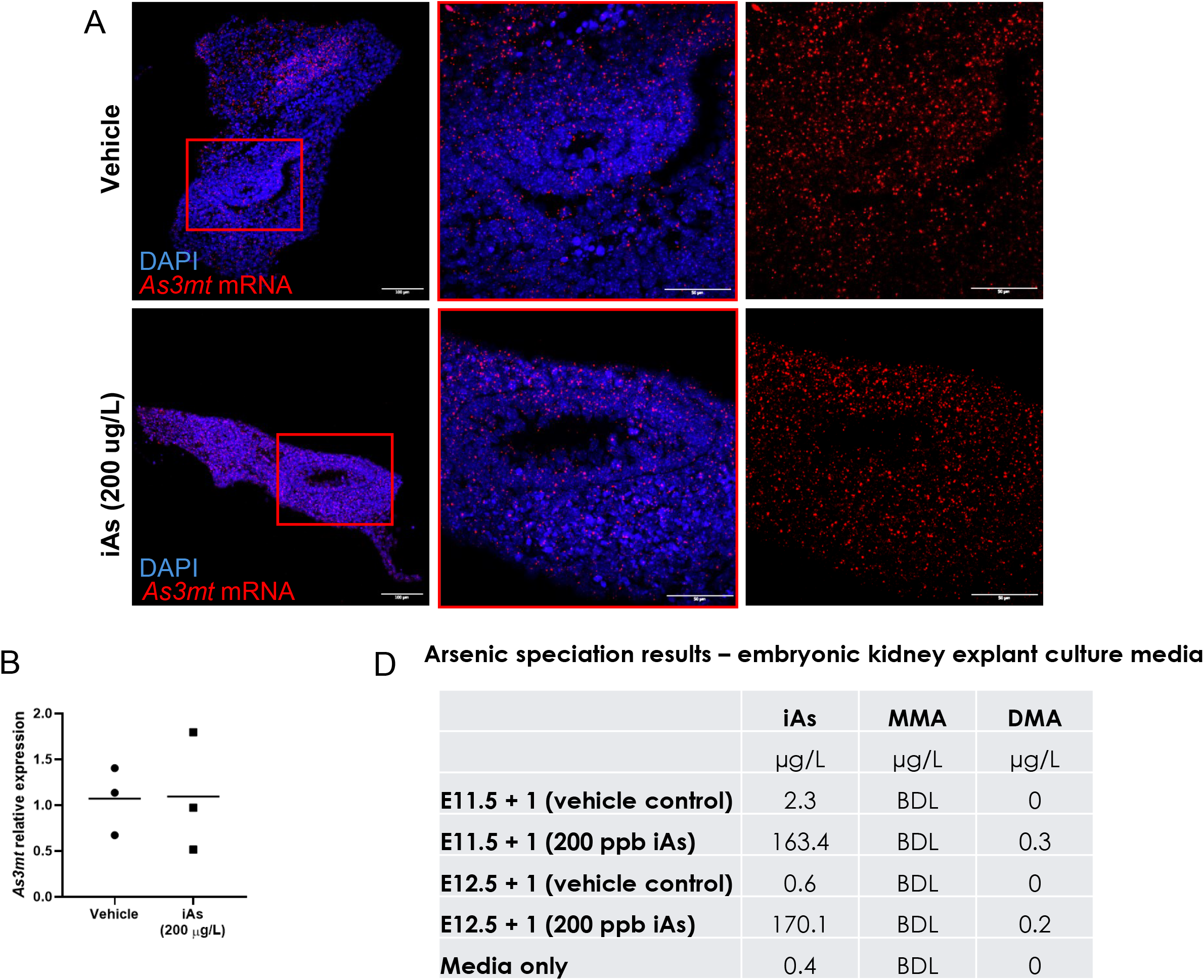
Embryonic mouse kidneys express As3mt and methylate iAsIII. (A) *As3mt* transcripts visualized *in situ* from E11.5 mouse kidneys cultured for 48 h in the presence or absence of 200 μg/L iAs. After culture, explants were fixed, embedded and sectioned to perform RNAScope against *As3mt*. (B) Nuclei were counterstained with DAPI (blue) Representative images are shown. Bar equals= 100 μm. n=3. (B) Relative gene expression of *As3mt* from RT-qPCR of E11.5 mouse kidneys cultured for 48 h in the presence or absence of 200 μg/L iAs. Data were analyzed using t-test. n=3. (C) Arsenic speciation results from the culture media from an E11.5 + 1 day and E12.5 + 1 day embryonic kidney explant culture experiment. The data represents the concentrations of iAs, MMA and DMA detected in the culture media by Liquid Chromatography Inductive Coupled Plasma Mass Spectrometry (n=1 experiment). BDL=below detection limit.

### Exposure to inorganic arsenic in utero affects nephron endowment

To determine if iAsIII has similar effects on nephrogenesis *in vivo*, pregnant female C57Bl/6J (wild-type) mice were exposed to 200 μg/L iAsIII in their drinking water beginning at embryonic day 0 and continuing throughout gestation and the weaning period. The offspring were euthanized at postnatal day 14, and body weights, kidney weights, and nephron number counts were obtained. Histological staining of control and iAsIII-treated wild-type mouse kidney sections with periodic acid-Schiff (PAS) revealed no overt differences in kidney morphology following iAsIII exposure (**Fig 5A,B**). Although body weights of wild-type mice exposed to iAsIII *in utero* were slightly smaller (control=6.4g, iAsIII=5.7g, p=0.018), there was also no significant difference in kidney weights between iAs-treated versus control offspring (control=0.53g, iAs=0.47g, p=0.16), even when kidney weight was normalized to body weight (**Fig. 5E, F**). Whole kidneys underwent acid maceration to obtain glomeruli counts as a proxy of nephron number. This revealed no difference in nephron number between iAsIII-treated versus control offspring (mean total nephron number/kidney; control=10,212, iAsIII=10,483 p=0.67) (**Fig. 5G**).

**Fig 5.**
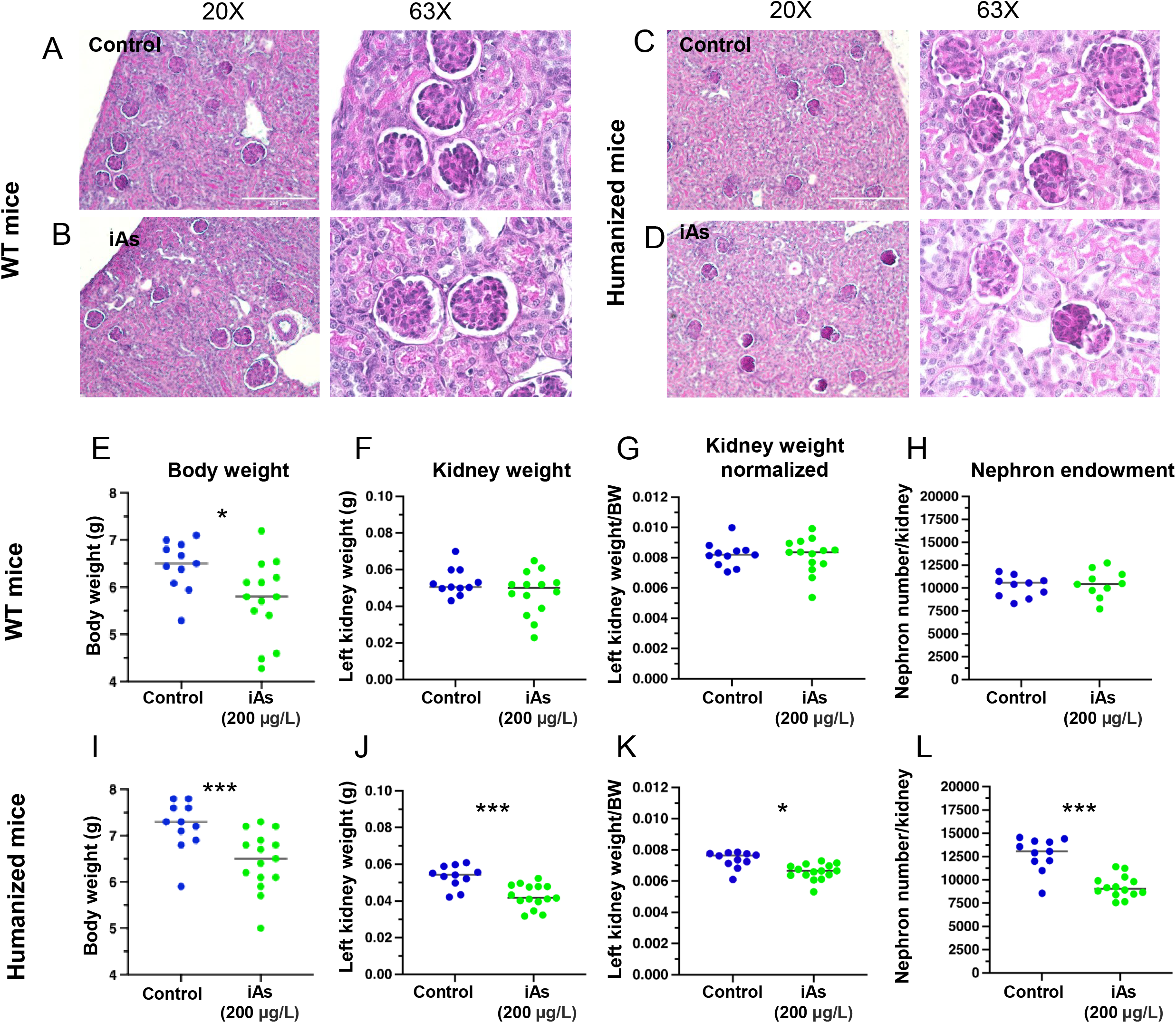
iAsIII exposure *in utero* decreases nephron endowment. Representative periodic acid-Schiff staining of left kidney sections of P14 wild-type (A,B) mice and humanized (C,D) mice from control conditions or iAsIII at 200 μg/L. Body weight, left kidney weight, kidney weight normalized to body weight (BW) and nephron numbers from of P14 wild-type and humanized mice from control and iAs exposed dams are shown (E-L). Each dot represents an individual mouse, and the horizontal lines indicate the mean. Data were analyzed with unpaired t-tests (parametric or non-parametric depending if data was normally distributed) *** = p < 0.0001; **=p<0.001; *=p<0.05. n=25 wild-type mice; n=26 humanized mice. Scale bar = 200 μm.

The absence of an effect on nephron number in wild-type mice could be due to rapid metabolism of iAsIII to DMA from the murine As3MT enzyme. To recapitulate human-like metabolism of iAs in a mouse model, a transgenic knock-in mouse harboring the human *AS3MT* gene was exposed to iAsIII (‘humanized mice’) ^17^. Homozygous humanized *AS3MT* breeding pairs were used to generate offspring. Pregnant homozygous females were exposed to 200 μg/L iAsIII in their drinking water or regular tap water throughout gestation and weaning. PAS staining of humanized mouse kidney sections revealed mild differences in glomeruli morphology in the iAs-treated humanized mice compared to control humanized mice – glomeruli appeared condensed and darker in colour, although this was not quantified (**Fig 5C, D**). Importantly, the kidney weights of iAs-exposed humanized offspring were significantly smaller compared to controls (control=0.53g, iAs=0.42g, p=0.0003), even when normalized to body weight (**Fig. 5H, I**). iAsIII-exposed humanized *AS3MT* pups also had a significant reduction in nephron number at P14 compared to tap-water exposed pups (control=12,773 nephrons/kidney, iAs=9,268 nephrons/kidney, p=0.0001) (**Fig. 5J**). Together, these data indicate that *in utero* exposure to iAs disrupts kidney growth and nephrogenesis *in vivo*. The spatial expression of *As3mt/AS3MT* was examined in both models using species-specific probes. In the wild-type mouse, *As3mt* was ubiquitously expressed and accentuated in glomerular structures. In contrast, in the humanized mouse, *AS3MT* appeared less abundant and little expression was seen in glomeruli (**Fig. 6A-C**). To investigate further, primers were designed to amplify both genes simultaneously by RT-qPCR (Table 1) and this revealed a decreased number of human *AS3MT* transcripts relative to mouse *As3mt* (**Fig. 6D**). This suggests there are regional differences in iAs methylation capacity based on As3MT genotype. No differences in either mouse line were noted when comparing enzyme expression between controls and iAsIII-exposed kidneys.

**Fig 6.**
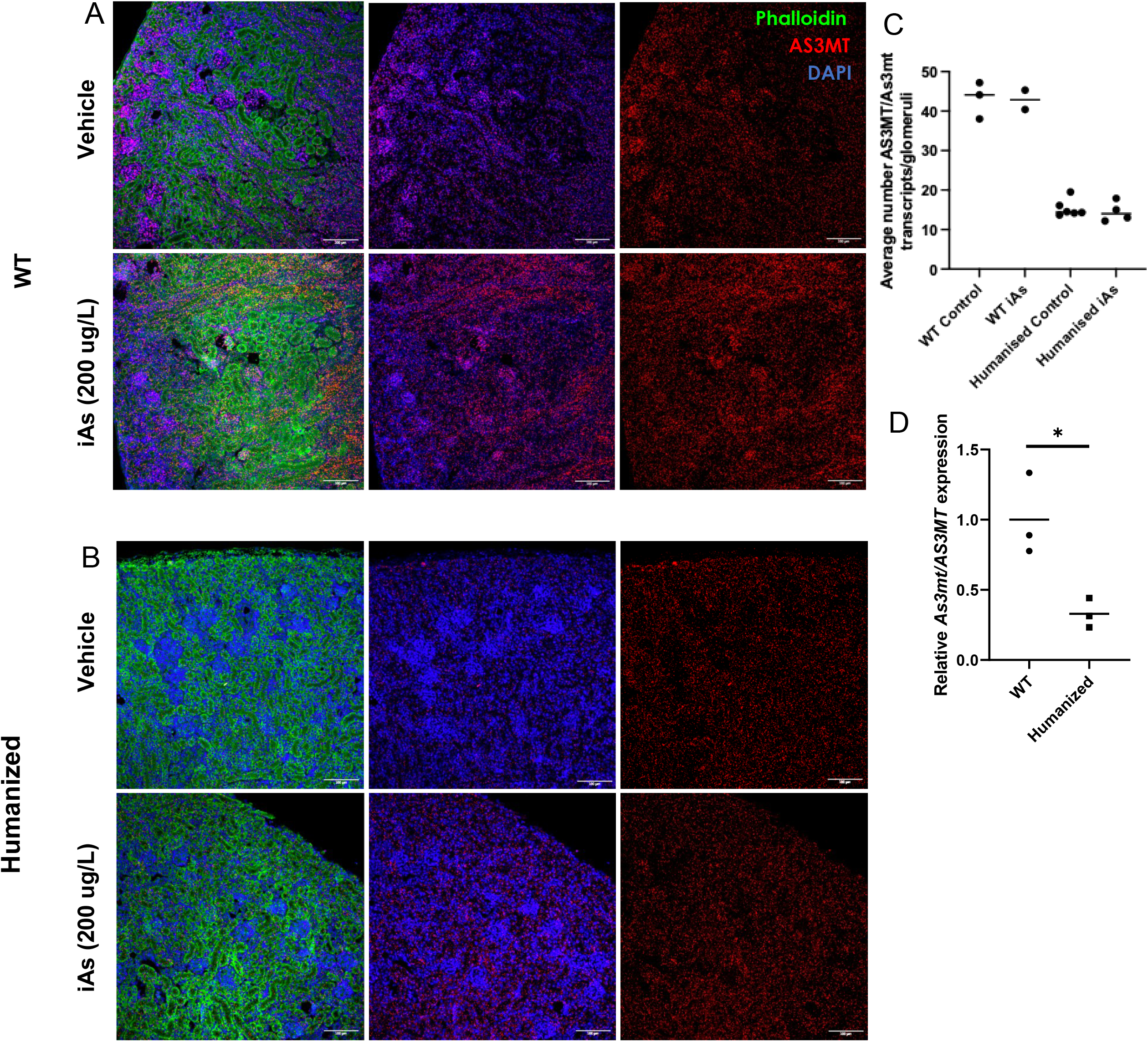
P14 mouse kidneys express As3mt/AS3MT. Wild-type (A) and humanized (B) dams were given iAsIII (200 μg/L) or tap water (vehicle) throughout gestation, and the pups were euthanized at P14 to retrieve and section the kidneys. As3mt/AS3MT expression was analyzed using RNAScope (red) and sections were counterstained with phalloidin (green) to reveal tubular structures and DAPI (blue) to visualize nuclei. Representative images are shown. Horizontal lines on the graph equals=mean. Glomeruli were delimited to create a region of interest and quantify the number of transcripts in wild-type (C) or humanized mice (D) Bar equals=mean: n=3/mice per group analyzed. Each dot represents the average number of As3mt/AS3MT transcripts per glomeruli based on the quantification of at least two images per mouse. Scale bar = 100 μm.

The *in vivo* data was stratified by sex, and no significant differences in kidney weights, nephron counts and As3mt/AS3MT mRNA expression were observed between female and male mice.

## Discussion

Our work demonstrates the negative impact of arsenical exposure on nephron endowment and kidney growth. iAsIII and the methylated metabolite, MMAIII, impair ureteric bud branching morphogenesis in mouse embryonic kidney explants, while MMAV and DMAV do not. Impaired ureteric bud branching is likely secondary to increased apoptosis of metanephric mesenchymal cells with diminished expression of GDNF. *In vivo*, a moderate to high dose arsenic exposure compromises nephron endowment in a model of humanized mice, whereas this same iAs concentration had no effect on wild-type mice. To our knowledge, this is the first report that effectively links *in utero* iAs exposure to decreased nephron endowment. This complements existing evidence that arsenic can interfere with organogenesis and normal embryo development ^23–26^.

Total nephron number varies greatly in humans ranging between 200,000 and 2.7 million nephrons and is influenced by genetics, sex, ethnicity, gestational length, and environmental factors ^27,28^. Low nephron endowment at birth predisposes individuals to CKD as part of normal aging with loss of nephrons and/or injury if diabetes or hypertension or other insults occur.

While the impact of perinatal exposures to toxicants on nephron endowment is less understood, growing evidence indicates that such early-life exposures could have lasting detrimental effects on kidney health. While the human studies of early-life exposure are limited, *in utero* and postnatal exposure to inorganic arsenic in Bangladesh, using maternal urinary As concentration during pregnancy and infant urinary As concentration, was associated with higher systolic and diastolic blood pressure in children at 4.5 years of age ^29^. The people of Antofagasta, Chile were exposed to a high concentration of iAs in their drinking water, 870 ug/L, between 1958-1970. Mortality rates were reported in Antofagasta compared to the rest of Chile in young adults (aged 30-40 years), revealed a higher standardized mortality ratio due to chronic kidney disease, suggesting a correlation between arsenic exposure and kidney health ^30^. Here, we propose that prenatal arsenical exposure is a risk factor for CKD in adulthood through attenuation of nephron endowment.

We report a critical embryonic development window of iAsIII susceptibility during early mouse nephrogenesis in our kidney explant model. Others have observed that differentiated tissues are less sensitive to arsenic toxicity i.e. epithelial cell populations are less affected by iAs than undifferentiated stromal cells in a model of gut organoids ^31^. At E13.5, mouse kidneys have a well-developed ureteric bud epithelium and the majority of the metanephric mesenchymal cells have already committed to a lineage. In contrast, at E11.5 the kidneys have a nascent epithelial T structure and a highly potent metanephric mesenchyme. Indeed, most of the apoptosis observed in iAsIII-exposed E11.5 embryos occurs in the metanephric mesenchyme: little cell death is observed in epithelialized ureteric buds and branches. The metanephric mesenchyme secretes GDNF that drives ureteric bud branching via signaling through the RET tyrosine kinase receptor expressed at the ureteric bud tips. The increased apoptosis in the metanephric mesenchyme in iAsIII-treated explants results in less production of *Gdnf* mRNA transcripts and secondarily impaired ureteric bud branching. Numerous genetic models, as well as environmental exposure models, have demonstrated a direct link between the depletion of metanephric mesenchymal populations and decreased ureteric bud branching, which results in hypoplastic kidneys with decreased nephron endowment ^10,32^. The increase in apoptosis is consistent with well-documented effects of arsenic in other biological systems in which it suppresses cell cycle checkpoint proteins, changes growth factor expression, inhibits DNA repair, alters DNA methylation, and increases oxidative stress ^33^.

Both iAsIII and MMAIII impaired ureteric bud branching and kidney growth in explants. RNAscope and the presence of DMA in the iAsIII-treated explants, even at a low concentration, confirmed that As3mt is functional, however it is likely that the enzymatic function of As3mt is overwhelmed by the concentration of iAsIII that was tested. We suspect that the explants may be depleted of co-factors that are necessary for both arsenic bio-transformation and DNA methylation, as observed by others ^34^. The relative amounts of glutathione and S-Adenosyl methionine in iAsIII-treated explants may be rate-limiting for maximal function of As3mt. *As3mt* was also detected by RNA-scope in P14 kidneys. While no difference in expression was detected between P14 kidneys when comparing control and iAsIII-exposed dams in the wild-type or the humanized mice, we did observe less *AS3MT* transcripts in the glomerular structures of humanized mice. This suggests that specific lineages within the kidney, like podocytes, may be more vulnerable to iAs exposure due to less enzyme activity. This is consistent with a previous characterization of this transgenic strain: As3mt expression in humanized mice is decreased in highly vascularized tissues compared to wild-type mice ^17^.

The inclusion of the *AS3MT* humanized mice in this study was crucial given the divergence of As3MT in mice and humans, which renders different methylation efficiency (and expression patterns) to these orthologues. Mouse and human As3MT share only ∼75% of primary amino acid sequence ^35^. As a result, the mouse enzyme is more efficient at methylating arsenic which causes mice to excrete around 90% of iAs from a single dose in 48 h, whereas in humans the biological half-time of iAs is 4 d ^36^. Thus, mice are markedly less susceptible than humans to the adverse effects of iAsIII exposure. Indeed, at 200 μg/L iAsIII, we detected a difference in the renal phenotype between wild-type and humanized mice. A decrease in nephron endowment, as well as smaller kidneys even after body weight normalization, highlights a specific effect of iAsIII on the developing kidneys. Importantly, others have shown that exposures up to 400 μg/L iAs for 4 weeks showed no signs of general toxicity (e.g., severe weight loss or gross pathology in tissues) in either male or female humanized mice ^17^. This suggests that the dose employed in the present study could be relevant for humans, especially in regions with suspected iAs contamination where no other signs of intoxication may be apparent. In this scenario, the impact of iAs could be rescued by supplementation with compounds like folate, selenium or zinc that can reportedly antagonize the mechanisms of arsenic toxicity ^37,38^.

In conclusion, our data shows for the first time that the developing kidney is vulnerable to arsenical-induced nephrotoxicity, suggesting that *in utero* exposure to arsenicals can jeopardize childhood and adult kidney health. Further research on the *AS3MT* humanized mice exposed to iAsIII and its metabolites *in utero* is necessary to establish a direct link between arsenical exposure during development and CKD.

## Supporting information

Suuplemental Fig.1-3

## Figure legends

**Supplemental Fig 1.** Low iAsIII concentration does not affect early ureteric bud branching or kidney growth. Embryonic kidneys were dissected from mouse embryos at E11.5 and cultured in the presence or absence of 20 μg/L iAsIII. Ureteric branching index (A) was calculated as (N_T48_-N_T0_)/N_T0_ from fluorescent images (B). Data were analyzed using paired t-tests. Kidney perimeter was measured at each timepoint from brightfield images (D). Data were analyzed using two-way ANOVA. Representative epifluorescence images (D) at T0, T24 and T48h of culture are shown. Scale bar equals 100 μm. n=18. Embryos are collected from at least 3 dams.

**Supplemental Fig. 2.** Representative bright field images of kidney explants shown in Figure 1 and Figure 2 are shown.

**Supplemental Fig. 3.** Six2 and WT1 expression pattern is not affected by iAsIII treatment. E11.5 kidneys were dissected and cultured for 48 h in the presence or absence of 200 μg/L iAsIII. Kidney explants were fixed/permeabilized, blocked and incubated with antibodies against Six2 (cyan) and WT1 (magenta) for whole mount immunofluorescence. Anti-rabbit and anti-mouse antibodies coupled to Alexa 488 and Alexa 555, respectively were used as secondary antibodies. Nuclei were counterstained with DAPI. Expression of both proteins is concentrated around the ureteric bud tips and does not appear different between samples. Bar equals: 100 μm. Images shown are maximum intensity projections of 5-6 stacks separated by 10 μm. NC=negative control, no primary antibody incubation.

