## Supplementary figures and images for "Mouse nephron formation is impaired by moderate-dose arsenical exposure"

### Suuplemental Fig.1-3

Supplemental Fig. 1

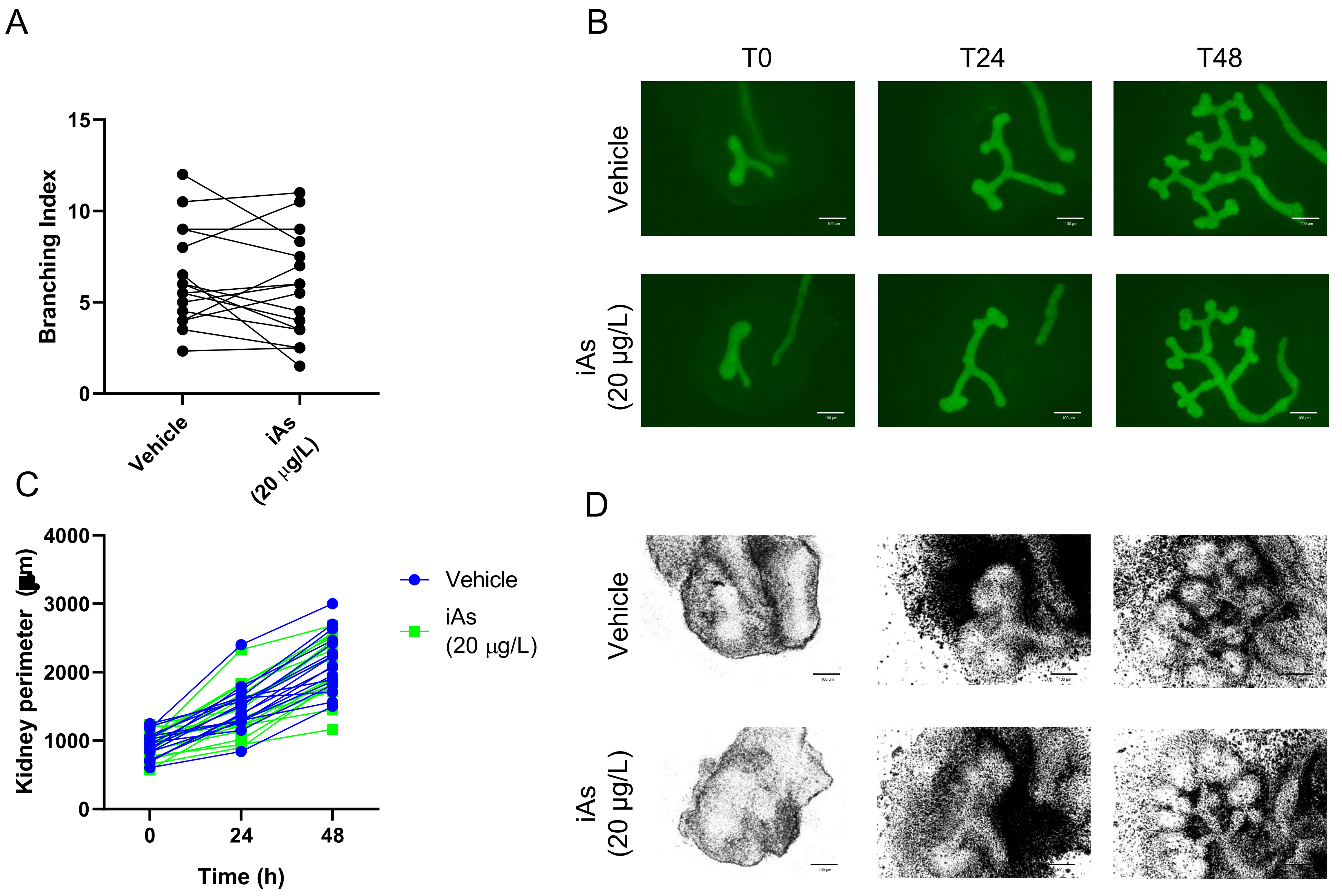

Supplemental Fig. 2

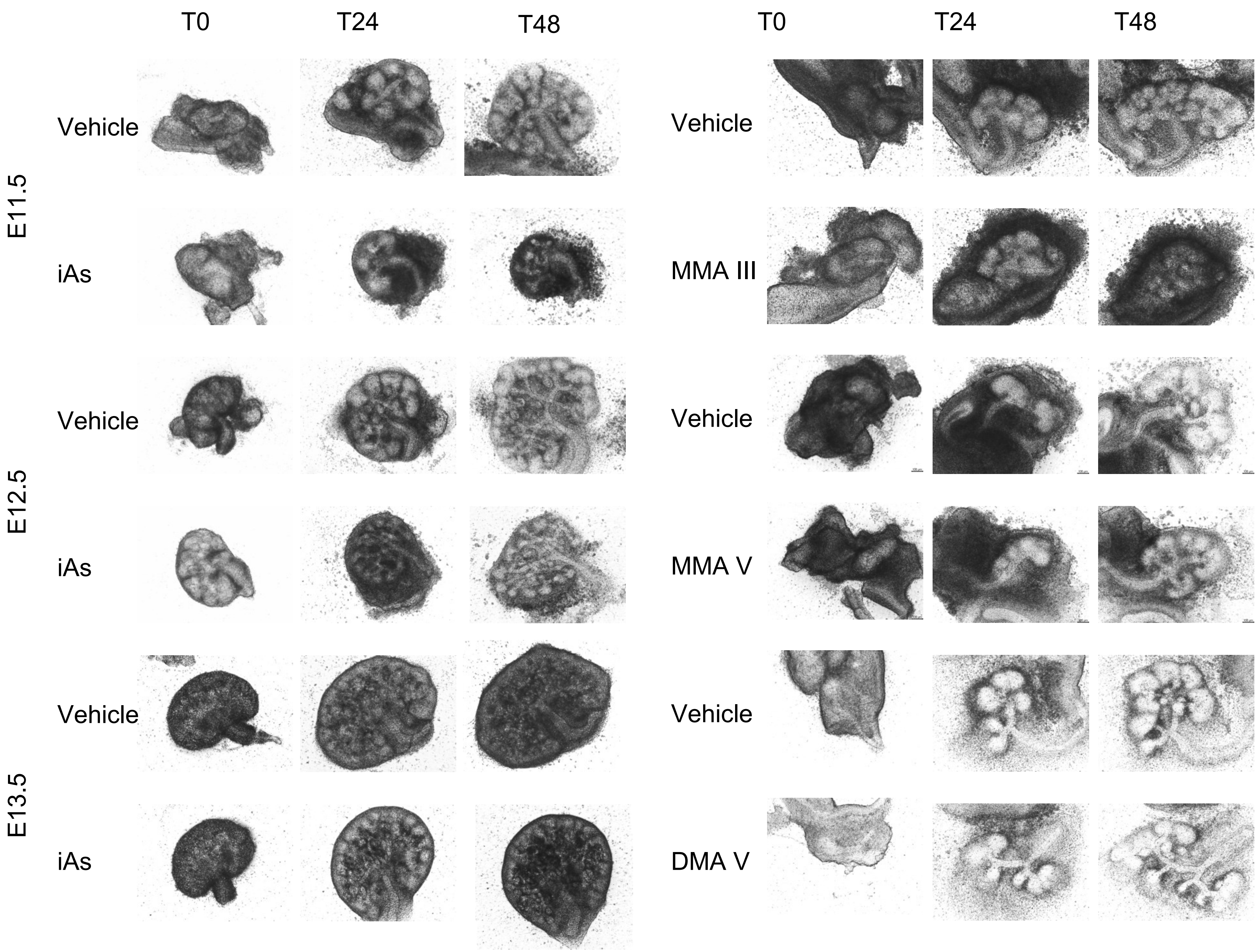

Supplemental Fig. 3

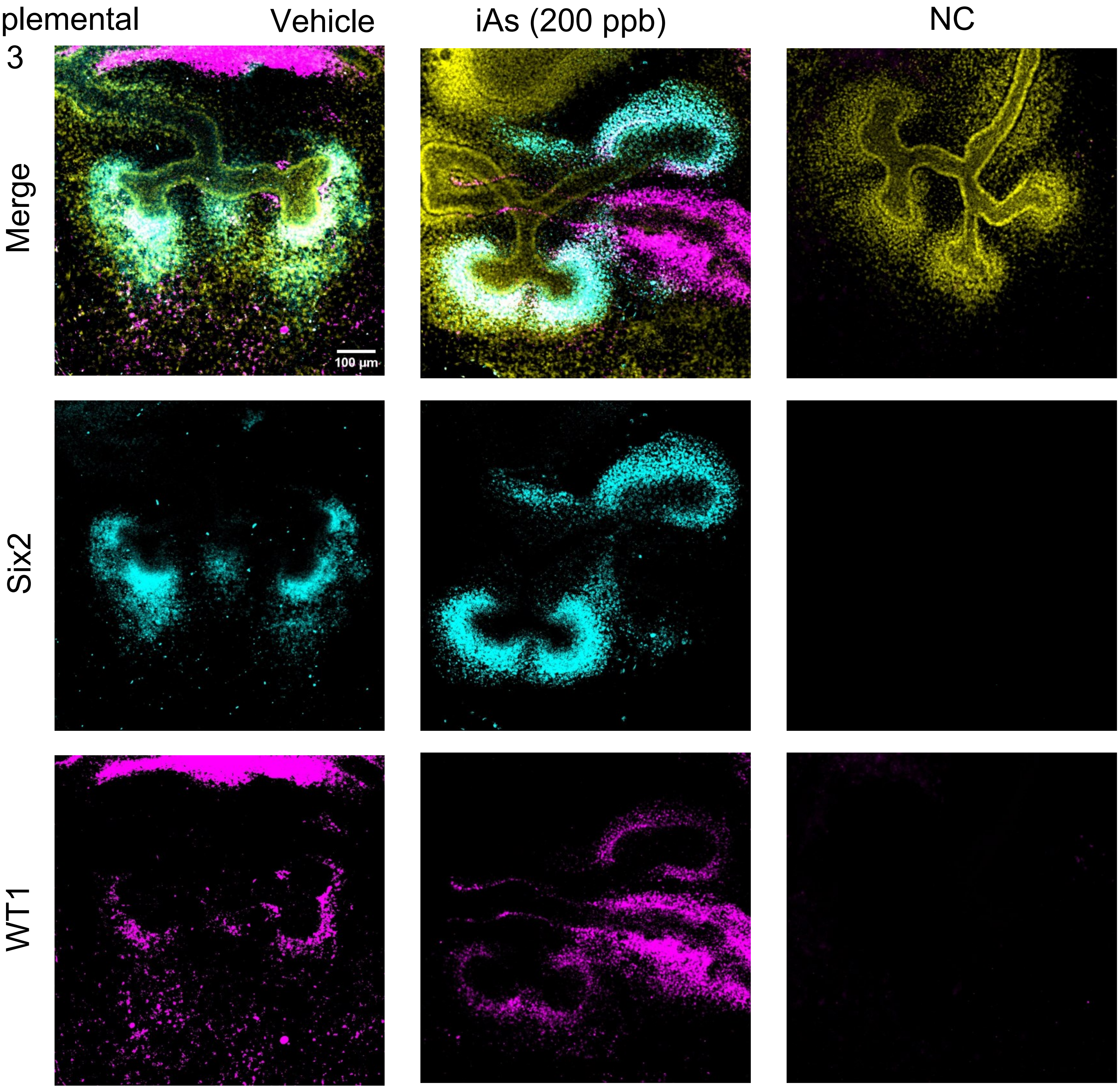
